# 200 years of river engineering: changes in meta-ecosystem resilience and connectivity

**DOI:** 10.64898/2026.07.03.736339

**Authors:** Johannes L. Kowal, Sarah M. Gross, Gertrud Haidvogl, Thomas Hein, Severin Hohensinner, Andrea Funk

**Affiliations:** Department of Ecosystem Management, Climate and Biodiversity, Christian Doppler Laboratory for Meta Ecosystem Dynamics in Riverine Landscapes, Institute of Hydrobiology and Aquatic Ecosystem Management, BOKU University, Vienna, Gregor-Mendel-Str. 33, DG 1180 Vienna, Austria

**Keywords:** Biodiversity, Networks, Functionality, Regulation

## Abstract

This study investigates changes in habitat connectivity and meta-ecosystem resilience between 1817 and 2022 along a 150 km section of a large European river (Danube) and its adjacent floodplains. The analysis was based on a time series of functional habitat networks (graphs) constructed from historical records and remote sensing data, integrating habitat suitability and dispersal modes of functional organism groups. The results indicate that overall habitat availability declined by about 50% since 1817, leading to the near-complete loss of functional connectivity among dynamic habitats by 1910. Less dynamic habitats persisted or expanded but became functionally less connected. Connectivity-based habitat classification further revealed four distinct functional connectivity clusters and the near-complete loss of an originally dominant, dynamic, and highly connected habitat type. Furthermore, the results indicated a fundamental loss of meta-ecosystem resilience. This development was reflected by increased spatial modularity among functionally similar habitats and by reduced layer dissimilarity and structural robustness in multilayer networks representing the spatial habitat structure and functional habitat connectivity across different functional organism groups.

**Significance:** River engineering is widely recognized for causing habitat loss and fragmentation, yet its long-term consequences for functional habitat connectivity and meta-ecosystem resilience remain poorly understood. This study illustrates how 200 years of river engineering reorganized a vast, dynamic, and highly interconnected river–floodplain system into a fragmented and functionally homogenized state. The findings empirically expose the true scale of functional degradation, offering a vital and novel historical perspective that underscores the urgent need for large-scale, process-based restoration.

## 1 Introduction

Rivers and their adjacent floodplains form highly dynamic and interconnected habitat networks that support a disproportionate share of global biodiversity (Balian et al., 2007). However, these ecosystems are undergoing rapid and persistent transformation, facing escalating pressures arising from dams (Kowal et al., 2025), flood protection (Kuiper et al., 2014), transport (Dittrich et al., 2026), water use (Sabater et al., 2018), pollution (Pinheiro et al., 2021), land-use changes, and climate-driven alterations (Comte et al., 2021). Among these pressures, hydromorphological alteration through river engineering has been particularly consequential. Channelization, levee construction, impoundment, and floodplain disconnection have not only caused habitat loss or degradation but have also fundamentally reorganized river–floodplain systems by altering the spatial configuration and connectivity of habitat networks that sustain ecological processes (Pander et al., 2018). This development has substantially contributed to the global biodiversity crisis (Sayer et al., 2025) and undermined ecosystem services that support human societies (Petsch et al., 2022). As a consequence, several major policies, such as the EU Nature Restoration Regulation (European Commission, 2024), the EU Water Framework Directive (European Commission, 2000), and the EU Biodiversity Strategy (European Commission, 2010), have been aimed at halting further degradation of freshwater ecosystems and partially restoring what has been lost. A major challenge in achieving these goals remains the methods used to assess ecological status, as shifted baselines and missing reference conditions hinder precise evaluation of the current state (Free et al., 2024). Furthermore, assessments are commonly conducted without consideration of the broader spatial context of individual sites or of changes in the overall resilience resulting from long-term developments. As a consequence, processes and trends on the meta-ecosystem level (Cid et al., 2021), such as seasonal migration or dynamics between subpopulations, are disregarded. However, given the extent to which many freshwater ecosystems have been systematically degraded by human activities, particularly since the onset of industrialization in the 19th century, it is crucial to base long-term conservation and restoration strategies on a holistic understanding spanning multiple spatial and temporal scales. Moreover, socioeconomic constraints are making it impossible to restore freshwater ecosystems to conditions similar to those in largely undisturbed systems in most cases (Funk et al., 2026). Instead, it has been stressed that ecosystem management in times of global change must be based on the coupling of the natural and social dimensions, treating freshwater and other habitats as socioecohydrological systems (Hein et al., 2021) using anthropocene baselines (Kopf et al., 2015). However, such a framing can only lead to long-term sustainability if key ecological processes are truly understood across scales ranging from small spatiotemporal scales to the meta-ecosystem level. Moreover, management decisions need to take into account the transformations these systems have undergone over time and the consequential effect on their ability to buffer disturbances.

Attempts to reconstruct the effects of river engineering on functional habitat connectivity and meta-ecosystem resilience across appropriate time scales are scarce. The Danube in eastern Austria, a mostly anabranching system with a historically extensive floodplain, offers such an opportunity, due to its exceptional historical documentation. Using this meta-ecosystem as an example, the specific objectives of the present study were to (1) quantify changes in habitat availability and type composition between 1817 and 2022. Furthermore, (2) to estimate functional connectivity for multiple organism groups using reach-and catchment-level indices, which were then used to (3) identify functional connectivity clusters and their spatial reorganization over time. The final objective was to (4) assess system-level resilience based on CC modularity, multilayer network robustness, and inter-layer dissimilarity.

## 2 Methods

### 2.1 Spatiotemporal framework

This study examines a central European fluvial system of the Danube, spanning a roughly 150 km stretch between the cities of Melk (Austria) and Bratislava (Slovakia), flowing from the eastern foothills of the Alps into the Pannonian Plain (figure 1). The area is characterized by a cold climate with warm summers (Köppen classification: Dfb) and an annual precipitation range of 400 to 1000 mm (GeoSphere Austria, 2024). While resistant hillslope geology moderately confines the upper section, the study area has historically featured an anabranching river system encompassed by several extensive, highly dynamic floodplains (Hohensinner et al., 2013a). The analysis spans seven key points in time between 1817 and 2022. The first observation in 1817 refers to the system before the onset of systematic regulation, while subsequent observations capture key phases of hydromorphological regulation and recent conservation efforts. By 1857, the system was still predominantly dynamic. However, initial regulation measures had commenced, leading to the regulation of the first sections until 1875, particularly in the vicinity of Vienna. The year 1910 marks the full implementation of measures regulating discharges up to the annual average in the entire study area. Until 1965, additional levees were constructed everywhere except south-east of Vienna, resulting in the disconnection of floodplains even during discharges above the annual average. The year 1996 marks the highest level of regulation, when an additional major levee, located a few hundred meters from the main channel, further disconnected the floodplain on the left bank (in flow direction) east of Vienna. Furthermore, three major dams were constructed along the main channel and at the upper end of the Danube Canal (Vienna), a major side channel. As a consequence, 42% of the main branch remains fully impounded. Measures to enhance fish passability were not implemented at the time, except for a fish migration aid constructed during the 1990s at the most downstream dam. Until 2022, several measures were implemented to restore, create, and reconnect lost habitats. These included, among others, reconnecting side channels, creating gravel bars, and constructing fish migration aids at the three previously impassable dams.

**Figure 1:**
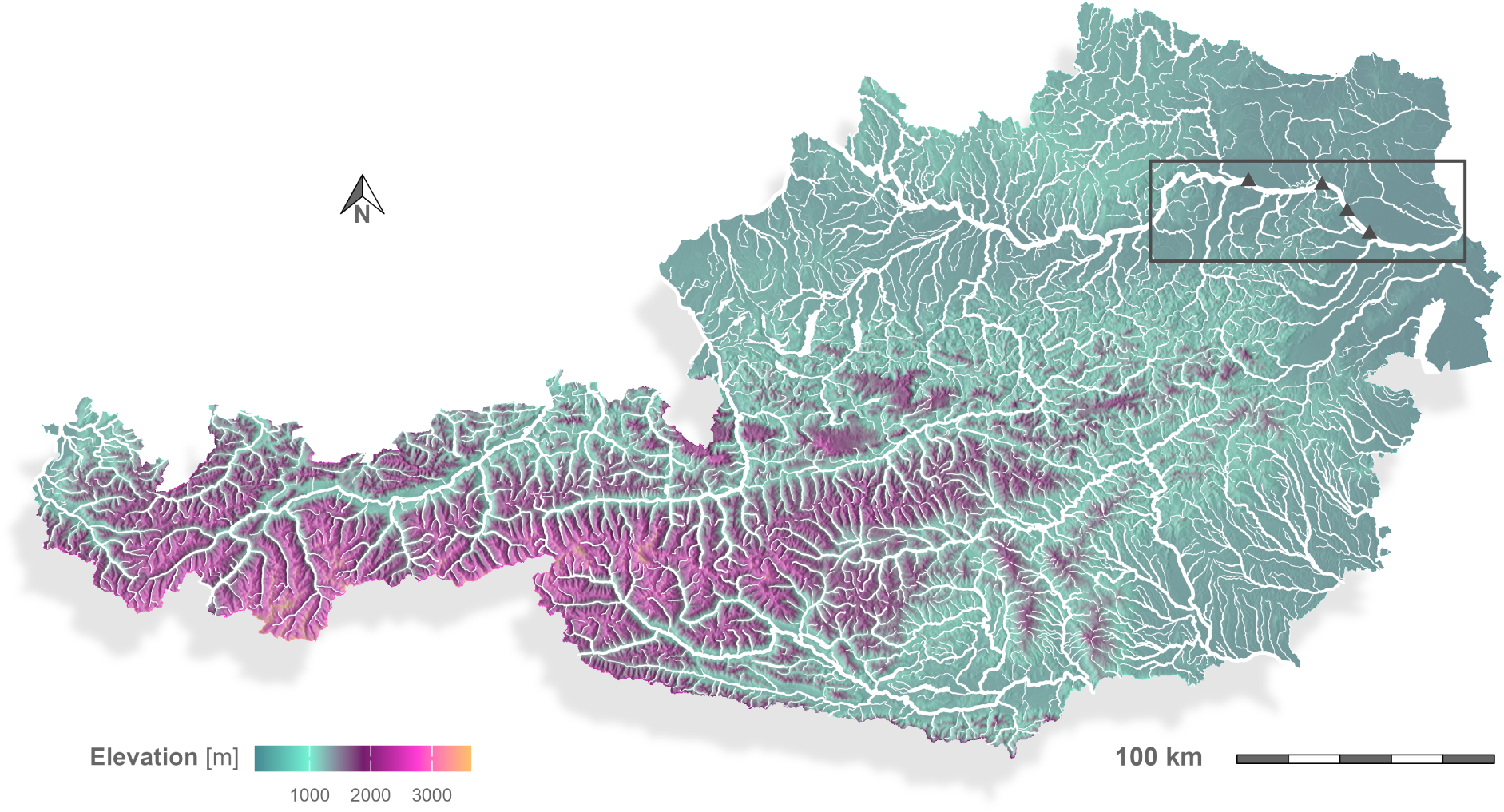
Visualization of the Austrian river network. Triangles indicate infrastructure complexes that are acting as transversal barriers and are located along the investigated section (bounding box) of the Danube. The predominant flow direction runs from west to east.

### 2.2 Reach mapping

River and floodplain waterbodies (hereafter referred to as reaches; figure 6) were mapped as spatial line features using remote sensing data for the most recent time point analyzed (2022). Earlier years were reconstructed from several historical sources (table 2; appendix) using a regressive, iterative approach proposed by Hohensinner et al. (2013b). Reaches connected to others were mapped as spatial line features from their upstream confluence(s) to their downstream confluence(s). Rather than the center of the river channel, the point of confluence was used as a connection point to map the length of individual reaches and thus generate a more accurate estimate of the provided habitat (figure 8; appendix). This caused the spatial line features, particularly those referencing wide reaches, to be technically disconnected. To generate a network (graph) for the calculation of functional connectivity indices (section 2.3), which correctly references all relevant connections between reaches, additional spatial line features (connectors) were added to bridge the described gaps and establish connections where appropriate.

In addition to the spatial mapping, reaches were further classified into seven habitat types *h* according to differences in their degree of hydrological connectivity to the main channel (figure 3B & 4). While generally following the classification framework proposed by Amoros et al. (1987) and later adapted by Hohensinner et al. (2011), these habitat types were additionally distinguished as impounded or non-impounded reaches.

**Figure 2:**
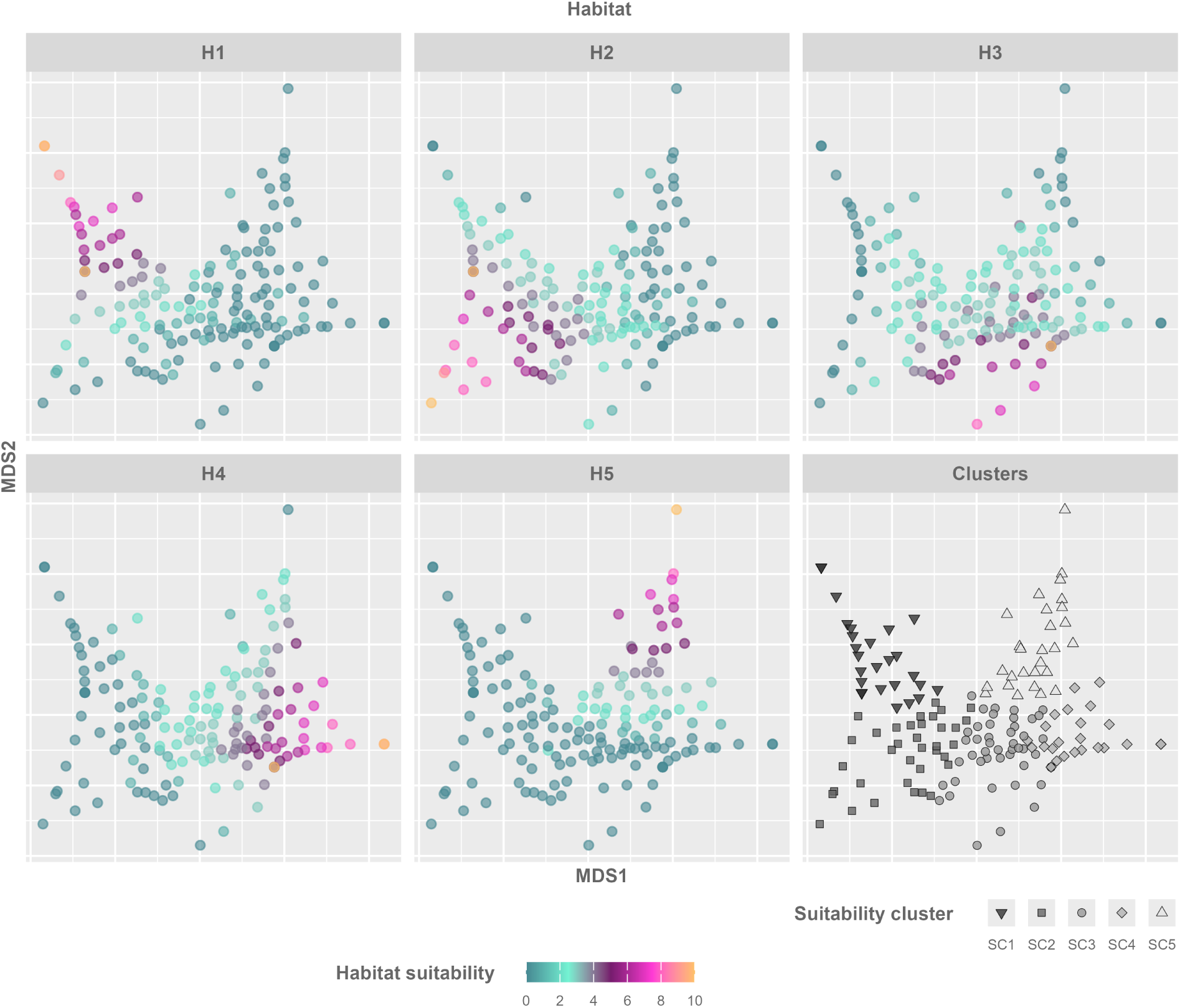
NMDS plots (*k* = 2; stress = 0.13; linear fit *R*^2^ = 0.95) visualizing the suitability of different habitats defined by Chovanec et al. (2005) for different aquatic and semi-aquatic taxa including fish, amphibians, macroinvertebrates, and macrophytes. The bottom right facet visualizes the grouping of taxa into suitability clusters (SCs).

**Figure 3:**
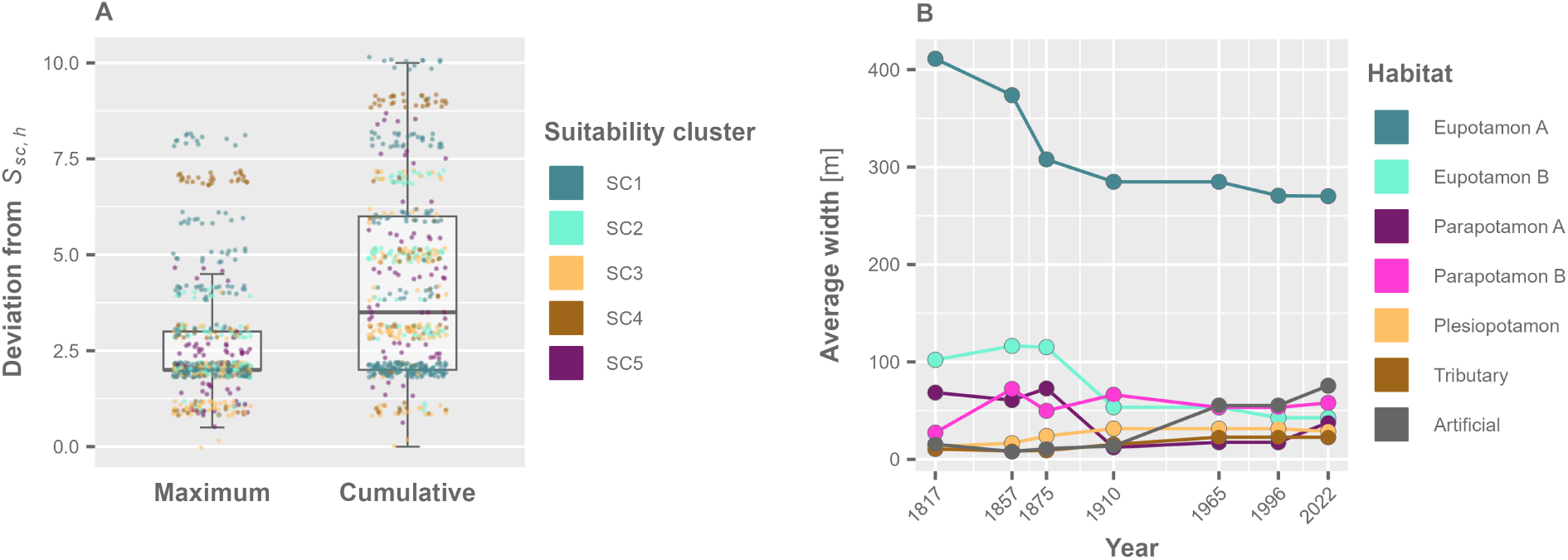
**A** Maximum and cumulative deviation of habitat suitabilities of taxa from *S_sc, h_*. **B** Average width of the river bed *W_y, h_* per habitat type *h* defined by Hohensinner et al. (2011) during different points in time *y*. Average widths of impounded habitats are not included.

**Figure 4:**
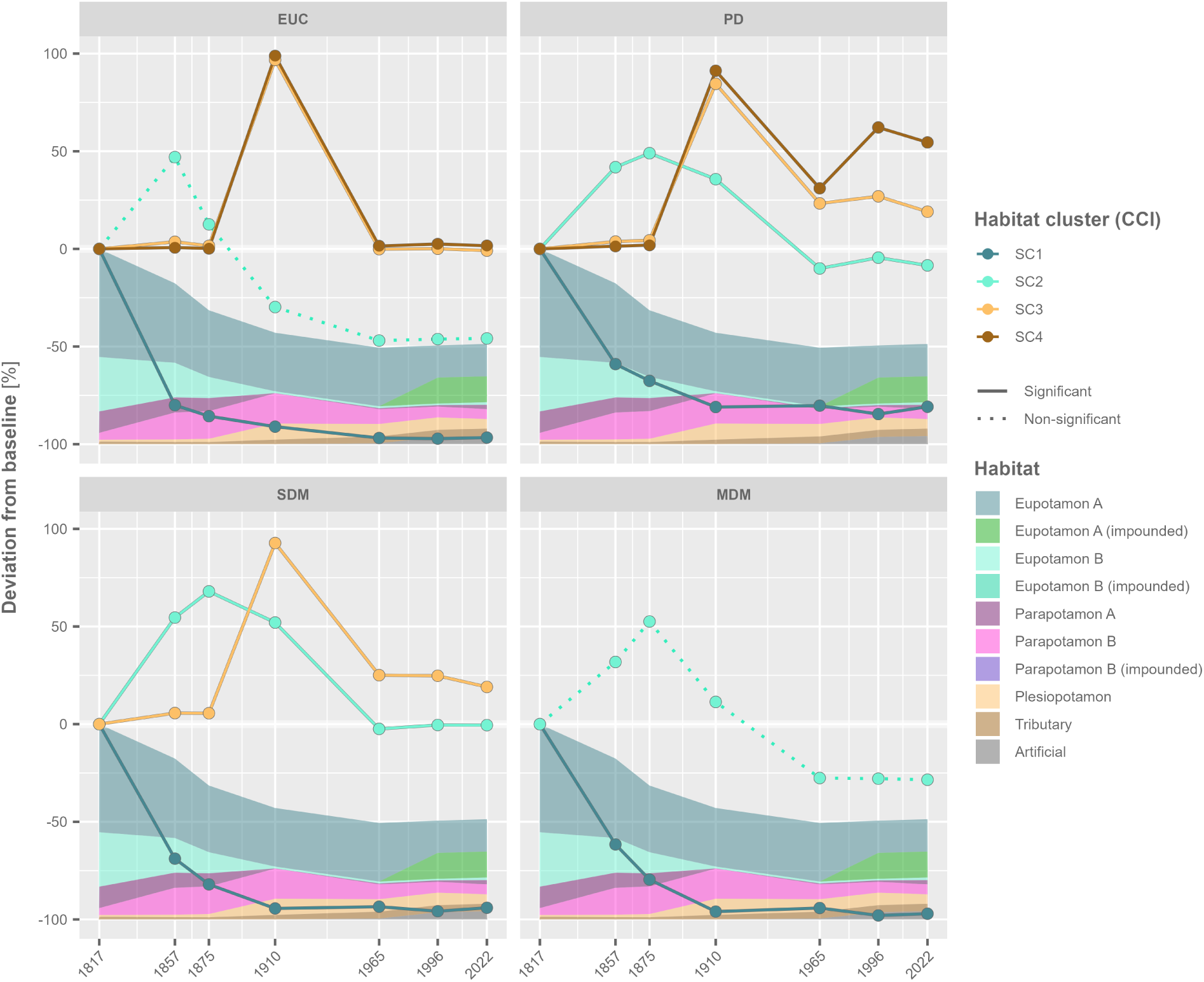
Catchment connectivity indices (CCI) and the cumulative area of habitats over time, visualized as the deviation from the baseline in 1817. CCI configurations in which the corresponding reach connectivity indices (RCIs) significantly explained the presence and absence of associated taxa are indicated by solid lines, whereas dotted lines denote non-significant configurations.

Finally, for every combination of habitat type and year, the average width of the river bed (figure 3B) was calculated by dividing the cumulative length of all spatial line features per habitat type by the cumulative area of the corresponding habitats. For this purpose, spatial line features were joined with corresponding habitat polygons from a dataset of the Viennese region, including the Lobau area (Hohensinner, 2008). This dataset covered approximately 18% of the analyzed stretch of the Danube in its current course. The distinction in average widths between years was deemed necessary because, over time, the widths of artificially disconnected side channels decrease due to siltation and other terrestrialization processes. The average width of a central, disconnected side channel of the Danube (locally referred to as the Old Danube) was calculated separately, as this habitat, classified as plesiopotamon (after disconnection), was substantially wider than other habitats of the same category.

### 2.3 Connectivity calculations

To assess functional connectivity from the perspective of aquatic and semi-aquatic floodplain taxa (figure 10; appendix) over time, changes in habitat quality, quantity, and connectivity were integrated through a network (graph) based approach. The following analysis was implemented in R (R Core Team, 2023) using packages from the tidyverse collection (Wickham et al., 2019) as well as riverconn (Baldan et al., 2022) and igraph (Antonov et al., 2023; Csárdi and Nepusz, 2006; Csárdi et al., 2026). The code used for the analysis is provided via the supplementary materials.

Based on the riverconn R package, a set of connectivity indices was defined based on combinations of habitat preferences and dispersal modes (table 1). For this purpose, a habitat network was constructed for each analyzed point in time (year *y*) based on the previously mapped reaches. While each reach generally defined a corresponding node in the network, long reaches were split into smaller segments *<* 500 [*m*] to ensure a spatial resolution detailed enough to investigate dispersal also on smaller spatial scales. The number of nodes in the resulting networks ranged between 7650 in 1817 and 4546 in 2022.

**Table 1:** Equations used for the calculation of connectivity indices.

| Definition | Equation |
| --- | --- |
| Weight of reach $j$ combining reach length $L_j$, average width $W_{y,h}$ and suitability $S_{sc,h}$. | $w_j = \ln(L_j W_{y,h}) \frac{S_{sc,h}}{10} \quad (1)$ |
| Weighted length (EUC, PD, SDM) of reach $j$ adjusted by habitat suitability $S_{sc,h}$ with $\varepsilon = 10^{-6}$ added to handle the case $S_{sc,h} = 0$ | $L_j^w = \frac{L_j}{\frac{S_{sc,h}}{10} + \varepsilon} \quad (2)$ |
| Weighted length (MDM) of reach $j$ adjusted by habitat suitability $S_{sc,h}$ with $\varepsilon = 10^{-6}$ added to handle the case $S_{sc,h} = 0$ | $L_j^w = \frac{L_j}{(\frac{S_{sc,h}}{10})^2 + \varepsilon} \quad (3)$ |
| Leptokurtic dispersal function (MDM, SDM) of the network distance $d_{ij}^{net}$ from reach $i$ to $j$ integrating over stationary and mobile subpopulations $P$. | $B_{ij}^{lep} = 2[0.5 P_{stat}(d_{ij}^{net}) + 0.5 P_{mob}(d_{ij}^{net})] \quad (4)$ |
| Exponential dispersal function, either with the euclidean distance $d_{ij}^{euc}$ (EUC) or the network distance $d_{ij}^{net}$ (PD) from reach $i$ to $j$ and the 5% dispersal threshold $T = 2000 m$. | $B_{ij}^{exp} = (0.05^{1/T})^{d_{ij}} \quad (5)$ |
| Effect of $k$ barriers on the path from reach $i$ to $j$ where $p_m$ is the upstream or downstream passability of barrier $m$ in the direction in which it is encountered. | $c_{ij} = \prod_{m=1}^k p_m \quad (6)$ |
| Network reach connectivity index of reach $i$, either with $B_{ij}^{lep}$ or $B_{ij}^{exp}$ in accordance with the dispersal mode assumed in the given configuration. | $RCI_i^{net} = \sum_{j=1}^n B_{ij} c_{ij} w_j \quad (7)$ |
| Euclidean reach connectivity index of reach $i$. | $RCI_i^{euc} = \sum_{j=1}^n B_{ij}^{exp} c_{ij} w_j \quad (8)$ |
| Network catchment connectivity index, either with $B_{ij}^{lep}$ or $B_{ij}^{exp}$ in accordance with the dispersal mode assumed in the given configuration. | $CCI^{net} = \sum_{i=1}^n \sum_{j=1}^n B_{ij} c_{ij} w_i w_j \quad (9)$ |
| Euclidean catchment connectivity index. | $CCI^{euc} = \sum_{i=1}^n \sum_{j=1}^n B_{ij} w_i w_j \quad (10)$ |

To reflect the importance of a reach *j* in a given configuration, weights *w_j_* were calculated (equation 1) based on the reach length *L_j_*[*m*], the average width *W_y, h_* [*m*] of the corresponding year *y* and habitat type *h* as well as a habitat suitability parameter *S_sc,h_*. Suitability parameters (table 3; appendix) were calculated based on habitat suitabilities of fish, amphibians, macroinvertebrates, and macrophytes taxa (Baart et al., 2013; Funk et al., 2017; Waringer et al., 2005). Habitat suitabilities from studies investigating organism groups other than macrophytes were set on a scale of 0 (low suitability) to 10 (high suitability) for five different habitat types (H1-H5) proposed by Chovanec et al. (2005). These habitat types were translated into the habitat types used to classify the previously mapped reaches, according to table 4 (appendix). For macrophyte habitat suitability values (Baart et al., 2013), habitat typologies were already aligned. However, this study only indicated the absence or presence of taxa in a given habitat, which was interpreted as a habitat suitability value of 0 or 10, respectively. From all taxa, distinct combinations of suitability values across the five habitat types were then grouped into five suitability clusters (SC1-SC5; *sc*) using the PAM algorithm (partitioning around medoids; Maechler et al., 2022). Suitability clusters were labeled after the corresponding habitat type where the highest suitability values occurred on average (figure 2). The number of suitability clusters was predefined based on the diminishing improvement in the within-cluster sum of squares with successive increases in the number of clusters (figure 9; appendix). Since the extracted suitability values were all on the same scale (0-10), no transformation was applied beforehand. Finally, suitability parameters *S_sc, h_*were calculated as the median of habitat suitability values per habitat *h* (see translation of habitat types; table 4; appendix) and suitability cluster *sc* with a standardization against the maximum possible value (10) added to all relevant formulas. The habitat suitability cluster SC5 and all associated taxa could not be further considered in the analysis, as it was not possible to map the corresponding core habitat H5 (paleopotamon) in a meaningful way across all investigated time points.

In addition to habitat suitability clusters, four dispersal modes were investigated. Fish were grouped into medium- (MDM; 30 *−* 300 [*km*]) and short-distance (SDM; *<* 30 [*km*]) migratory species according to Waidbacher and Haidvogl (1998). For these taxa, dispersal was defined through a leptokurtic dispersal kernel (equation 4) based on the shortest path 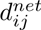 along the network from reach *i* to *j*. Leptokurtic dispersal was defined via two cumulative density functions of the normal distribution, reflecting the stationary *P_stat_* (50%) and the mobile *P_mob_* part of the population (50%). In the case of MDM taxa, cumulative density functions were configured based on the findings from Steinmann et al. (1937) as proposed by Kowal et al. (2025) using a mean of 0 and a standard deviation of 2600 [*m*] and 74200 [*m*] for *P_stat_* and *P_mob_*, respectively. For SDM taxa, the same standard deviations were applied after being scaled down by one order of magnitude in accordance with the classification of migration distances (Waidbacher and Haidvogl, 1998). While SDM taxa included mainly fish, taxa dispersing as fish parasites (several mollusks) were also assigned to this group. Furthermore, an exponential dispersal kernel (equation 5) was applied in the case of taxa classified as passive dispersers (PD) and those dispersing increasingly along Euclidean distances 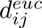 (EUC) and thus more independently of the network structure. Dispersal based on Euclidean distances was assumed for flying insects and amphibians, while passive dispersal in the downstream direction of network paths was assumed for all other remaining macroinvertebrate taxa and macrophytes. In both cases the dispersal threshold *T* was set to 2000 [*m*] so that 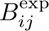= 0.05 when *d_ij_* = 2000 [*m*]. While true lifetime dispersal distances vary strongly across the considered taxa, a 5% threshold of 2000 [*m*] was considered to be a meaningful average (De Jager et al., 2018; Martínez-Gil et al., 2023; Ptatscheck et al., 2020).

Habitat suitability *S_sc h_* was further used to integrate higher or lower dispersal probabilities across reaches classified as habitats with higher or lower suitability, respectively. For this purpose, distances along the network 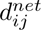 were calculated as the sum of weighted reach lengths *L^w^*of all reaches located along the shortest path from reach *i* to *j*. *S_sc h_* was integrated into *L^w^* through an inverse effect of *S_sc, h_* (equation 2). In the case of MDM taxa (equation 3), a quadratic inverse effect was used, which improved the accuracy as indicated by the validation with contemporary biotic data (section 2.4). Regarding EUC taxa, 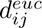 was calculated as the unweighted euclidean distance between reach *i* and *j*.

The multiplicative effect of all *k* barriers along the network path from reach *i* to *j* was integrated through upstream and downstream passability parameters *p_m_*, depending on the direction in which the barrier *m* was encountered (equation 6). For barriers without a fish migration aid, *p* was set to 0.3 downstream and 0 upstream. In the case where a barrier had been equipped with a fish migration aid, a value of 0.6 was used in the downstream direction, and 0.3 in the upstream direction. These passability values were based on the review from Noonan et al. (2011) and have already been applied in a previous study by Kowal et al. (2025).

Finally, spatially explicit reach connectivity indices (RCIs; equations 8 & 7) and spatially implicit catchment connectivity indices (CCIs; equations 10 & 9) were calculated. RCIs were further used to group reaches into four functional connectivity clusters (CC1-CC4; figure 6). For this purpose, highly correlated RCI configurations (Kendall’s *τ* > 0.7) were excluded. A z-transformation followed by an *ln*(*x* + 2) transformation was then applied, and the seven remaining RCIs (figure 11; appendix) were clustered across all analyzed points in time, applying the same algorithm used to generate suitability clusters. The number of clusters was again predefined based on the diminishing improvement in the within-cluster sum of squares with successive increases in the number of clusters (figure 9; appendix).

### 2.4 Validation

The approach was validated by comparing RCI configuration values (suitability cluster and dispersal mode) between sites with confirmed presence or absence of the corresponding taxa. While presences were indicated for sites where taxa had been recorded during sampling (Figure 11; appendix), absences were specified if the same sampling method had been applied, but the relevant taxa had not been recorded. The validation was performed on a pooled dataset containing observations of taxa across all organism groups considered in the definition of habitat suitability clusters. Data on fish, amphibians, odonates, and other selected macroinvertebrate taxa were used from Funk et al. (2020), Funk et al. (2013), and Chovanec et al. (2005). In addition, data on a broad range of macroinvertebrate taxa (Funk et al., 2017; Graf et al., 2013; Recinos Brizuela et al., 2025) as well as mollusks (Reckendorfer et al., 2006) and odonates (Reckendorfer et al., 2006) in particular, were added. Data on macrophytes originated from Baart et al. (2013). Observational data were unavailable (1817-1965) or scarce (1996) for the points in time analyzed, except for the most recent (2022). Hence, the validation was conducted based on RCI values representative of the system in 2022. In addition, observations were filtered to remove taxa that were less well represented by the general habitat suitability clusters (maximum deviation *>* 2 and cumulative deviation *>* 5 from *S_sc, h_*; figure 3A). Taxa for which observational data were available, and that were linked to the suitability clusters SC1-SC4 (SC5 was excluded since the corresponding core habitat H5 was not mapped), comprised fish and macroinvertebrates covering all four dispersal modes. Finally, a one-sided and unpaired Wilcoxon rank sum test was performed to assess whether RCI values of sites linked to presences were significantly higher (*p <* 0.05) compared to sites linked to absences. Tests were performed for each RCI configuration, defined by combinations of suitability clusters and dispersal modes, using a subset of the data downsampled to yield equal numbers of observations for presences and absences. Across all configurations, a clear trend toward higher RCI values at sites linked to presences was evident. These trends were statistically significant in all cases except for EUC taxa and MDM taxa linked to SC2, clearly underscoring the validity of the approach.

### 2.5 Resilience

Changes in the system’s resilience over time were inferred by assessing changes in CC modularity and through a multilayer network framework in which layers shared the same topology but differed in node (RCI) and link weights (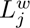) to reflect the functional connectivity of individual functional groups. Linking corresponding nodes across layers enabled resilience to be assessed simultaneously from multiple functional perspectives while retaining group-specific patterns. Resilience was then estimated based on network robustness and layer dissimilarity. To generate the link weights used in the multilayer framework, weighted reach lengths (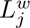), originally defined at the node level, were transferred to links. This was achieved by computing the arithmetic mean of 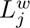 from the source and target nodes. The resulting weights were then scaled by the layer-specific maximum across years. In the case of inter-layer links, weights were defined uniformly as the arithmetic mean of intra-layer links within years. In addition, RCIs, scaled by the layer-specific maximum across years, were used to reflect the relative importance of individual nodes.

Robustness was evaluated by incrementally removing 1000 nodes with increasing functional connectivity (as measured by scaled RCIs) from the multilayer network. Subsequently, weighted shortest-path lengths were computed between a random subset of source nodes and all remaining nodes directly or indirectly connected to them. Each removal step was repeated 100 times, with 100 randomly selected source nodes per replicate, to account for stochasticity in node selection. This workflow was completed for each year’s multilayer network. As a measure of network robustness, the arithmetic mean of weighted path lengths was used, adjusted by the maximum possible number of paths to account for reachability.

Network dissimilarity between functional network layers was assessed as the cumulative Euclidean distance between scaled RCI vectors of corresponding nodes within each year’s multilayer network. This workflow produced a pairwise dissimilarity matrix for each year, indicating dissimilarities between RCI calibrations (layers). A decrease in functional dissimilarity between layers was interpreted as a loss in functional diversity and thus as an indication of lower system-level resilience.

Lastly, the modularity (Clauset et al., 2004) between functionally similar reaches was calculated using CCs as communities, with connector nodes (section 2.2, figure 8; appendix) grouped into a single additional cluster. Because CCs already represent a functional aggregation of RCIs, modularity was calculated outside the multilayer framework, using a single-layer network for each year. An increase in spatial modularization of functionally different parts of the investigated system was interpreted as functional isolation and thus as an indication of lower system-level resilience.

## 3 Results

### 3.1 Connectivity

The hydromorphological regulation of the investigated system resulted in a drastic loss and a shift in available habitat types (figure 4) with an overall loss of aquatic habitats in 2022 close to 50% relative to the 1817 reference. The most predominant changes occurred before 1910, characterized by a shift away from dynamic to less dynamic habitat types. After 1910, the most significant change in the composition of available habitat types was the impoundment of large sections along the main channel (eupotamon a), which peaked in 1996, further degrading dynamic habitats. Moreover, less dynamic habitat types, namely parapotamon B and plesiopotamon, indicated a continuous decrease after 1910. However, while the extent of parapotamon B habitat in 2022 was similar to that of 1817, it was still manyfold greater for plesiopotamon habitats.

When considering the functional connectivity of habitats (CCI; figure 4) alongside changes in availability and quality, a more differentiated picture emerged. Already in 1910, functional connectivity from the perspective of taxa adapted to highly dynamic conditions (SC1), which included a large share of the fish species native to the investigated system, was almost completely lost for short-distance migrators (SDM), medium-distance migrators (MDM), and organisms dispersing more independently of the habitat network structure (EUC). In the case of passive dispersers (PD), functional connectivity had decreased by approximately 75%. Regarding taxa associated with moderately dynamic habitats (SC2), an increase was observed until 1875 (1857 for EUC dispersers), followed by a decrease until 1965, after which it stagnated at a level similar to or lower than that of 1817. In the case of SC3 (less dynamic) and SC4 (isolated habitats), functional connectivity increased from 1875 to 1910, then declined sharply, resulting in final habitat accessibility levels similar to or higher than those in 1817, depending on the dispersal mode. Notably, the final levels of SC3 and, in particular, SC4 functional connectivity did not reflect the amount of plesiopotamon habitats, which are of central importance for both habitat clusters, and were manyfold higher in 2022 than in 1817. Hence, the results indicate that functional connectivity decreased with increased availability of these habitats.

Overall, reach connectivity indices (RCIs) indicated four distinct CCs (figure 5) that describe different functional habitat types, characterized by habitat quality and position within the habitat network, while accounting for different dispersal modes. CC1 was characterized by the highest average SC1 RCIs, high SC2 RCIs in MDM taxa, and the highest global average. CC1 was characteristic of the former anabranching, highly dynamic, and strongly connected meta-ecosystems. In contrast, while the global average remained high, CC2 indicated substantially decreased SC1 RCIs, more pronounced SC2 RCIs, and the emergence of SC3 RCIs. Therefore, CC2 is characterized by broader functionality and, consequently, a wider ecological niche that supports more generalistic taxa. Specifically, CC2 covered, e.g., side channels with a permanent connection only at the downstream end, as well as impounded reaches of the main channel. Furthermore, a third cluster, CC3, was very similar to CC1 in terms of relative RCI composition. However, the global average of RCIs was substantially lower, so that CC3 describes habitats such as tributaries and side channels within an overall simplified river-floodplain network, or Danube reaches that are disconnected from the floodplain but not impounded. Hence, CC3 indicates higher isolation than CC1 and less suitable conditions, particularly for SC1 but also for SC2 taxa. Lastly, CC4 was clearly dominated by SC4 and SC3 RCIs, with the global RCI average being the lowest of all CCs. Thus, CC4 describes non-dynamic habitats that are more isolated within the habitat network.

**Figure 5:**
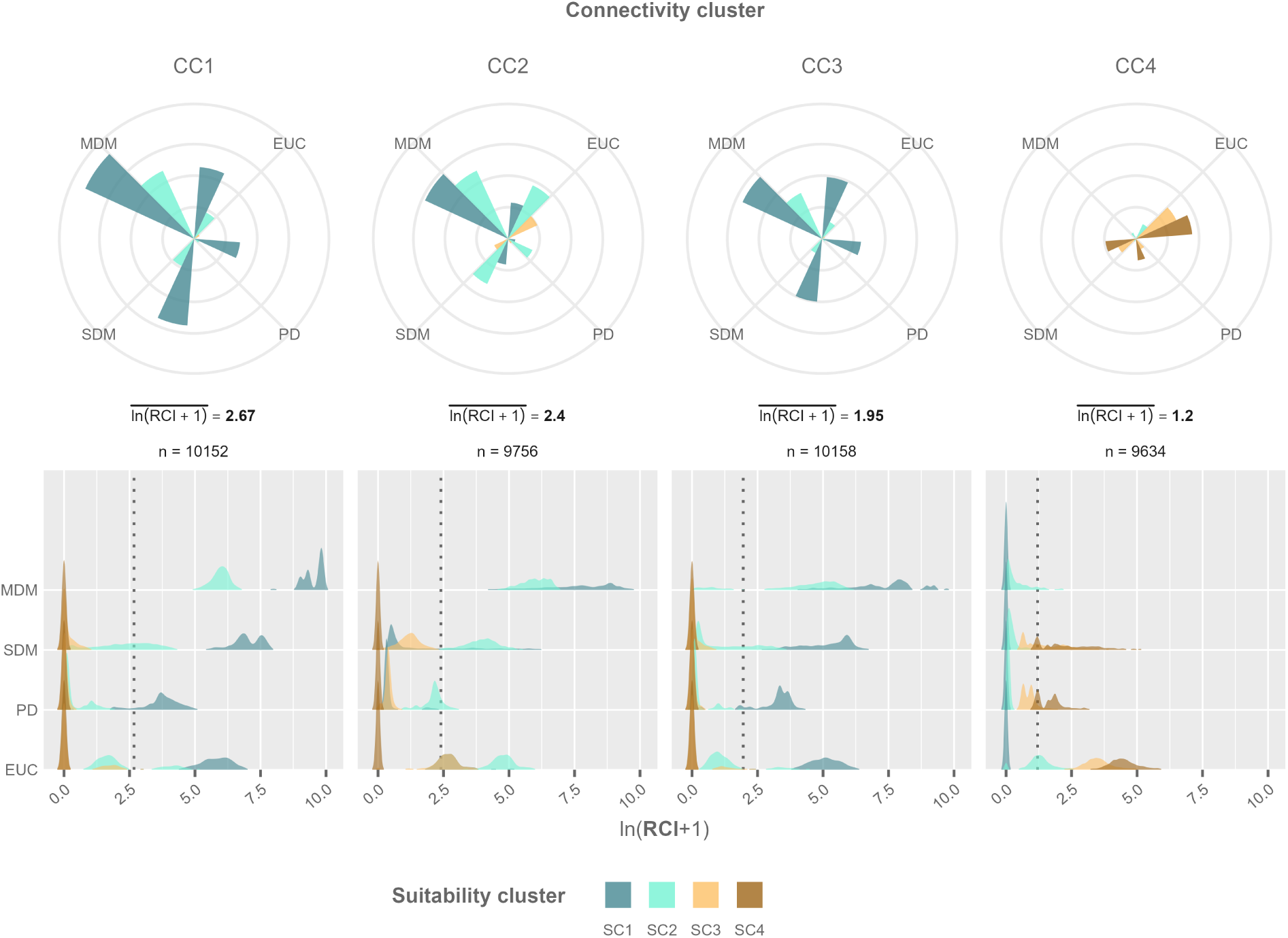
Circular barplots visualize the arithmetic mean of *ln*(*x* + 1) transformed reach connectivity indices (RCIs). The global arithmetic mean per connectivity cluster (CC) and the number of reaches across years (*n*) are indicated underneath. Density plots visualize the distribution of *ln*(*x* + 1) transformed RCIs, where vertical dotted lines indicate the arithmetic mean per CC. The color scale indicates suitability clusters (SC), while the labels MDM, SDM, PD, and EUC refer to medium-distance migrators, short-distance migrators, passive dispersers, and organisms dispersing more independently of the network structure, respectively.

**Figure 6:**
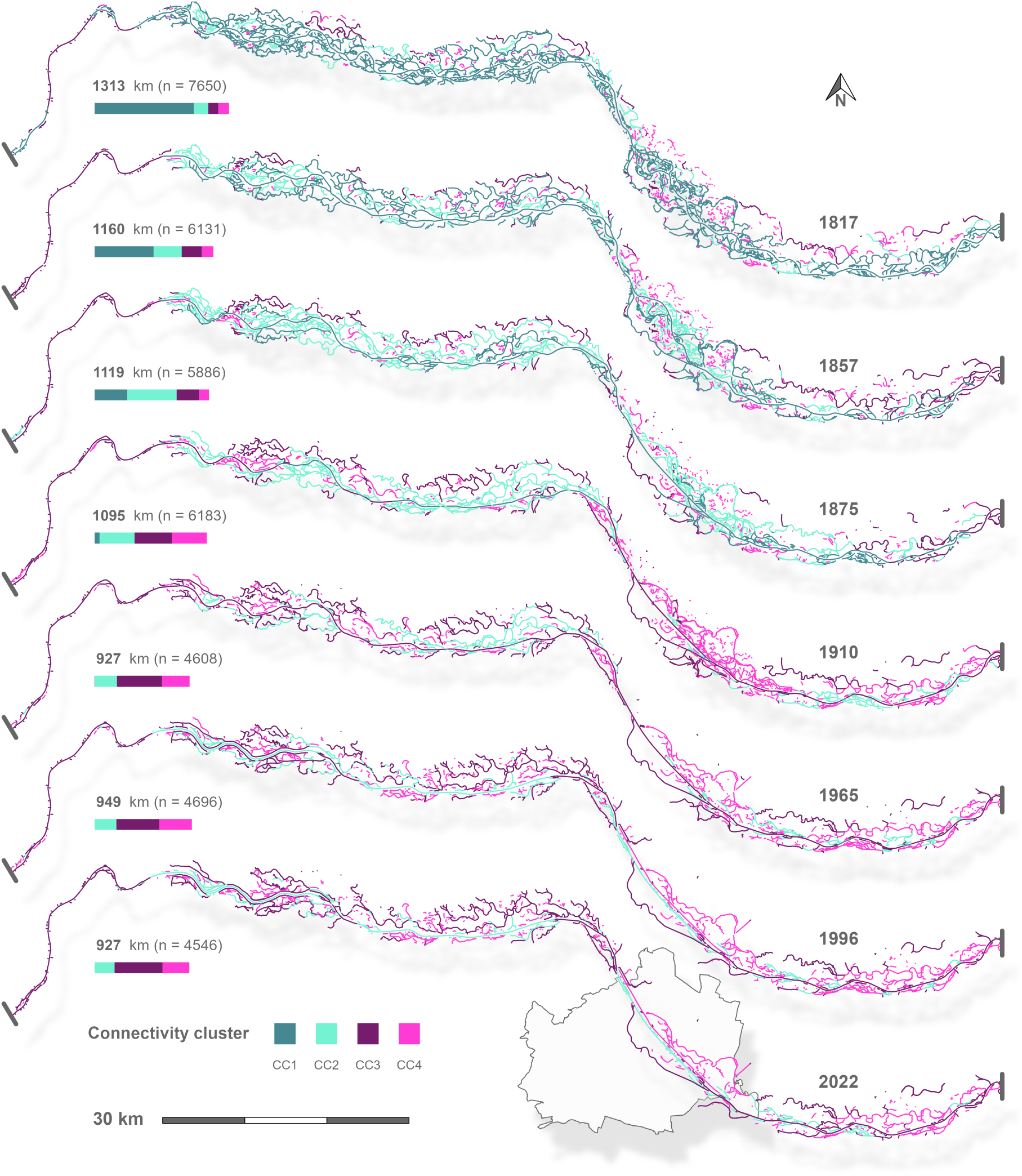
River network structure and connectivity clusters (CC) over time. The year is indicated on the right, while the total length of all reaches (bold) and the number of nodes (n) is indicated on the left. Horizontal bars indicated the total length of reaches per CC. The predominant flow direction is from west to east. For the 2022 river network, the city boundaries of Vienna are indicated (City of Vienna, 2025).

Over time, the extent and the spatial arrangement of CCs showed a dramatic shift (figure 6). While in 1817 CC1 clearly dominated, from 1857 to 1875 a substantial shift towards CC2, and to a lesser extent towards CC3, was observed. In addition, substantial losses of reaches occurred in the area surrounding the city of Vienna, thereby simplifying the network structure. Until 1910, a drastic shift from CC1 and CC2 to CC3 and CC4 occurred, particularly in the downstream half of the system. This trend continued over the next two time steps, 1965 and 1996, and was accompanied by the progressive loss of CC4 reaches. Notably, in 1965, almost all reaches previously classified as CC1 had shifted to CC2 or CC3, indicating that the most common functional habitat of the 1817 reference (historic pre-industrial condition) had effectively disappeared. After 1996, the system’s transformation had stalled, indicating that restoration measures produced only minor changes compared to the previous development.

### 3.2 System resilience

With time, and thus an increasing degree of river engineering, multilayer networks responded much more sensitively to the removal of nodes, indicating an overall reduced robustness (figure 7A). Collapse thresholds, at which the main network component became disconnected, were reached earlier (as indicated by a sudden increase in reachability-adjusted path lengths). In addition, the rate at which path lengths increased with the number of removed nodes became steeper. The result further indicated two main periods of change, the first between 1817 and 1857, and the second between 1910 and 1965. Notably, only minor differences were observed between 1996 and 2022, despite the implementation of restoration measures by 2022.

**Figure 7:**
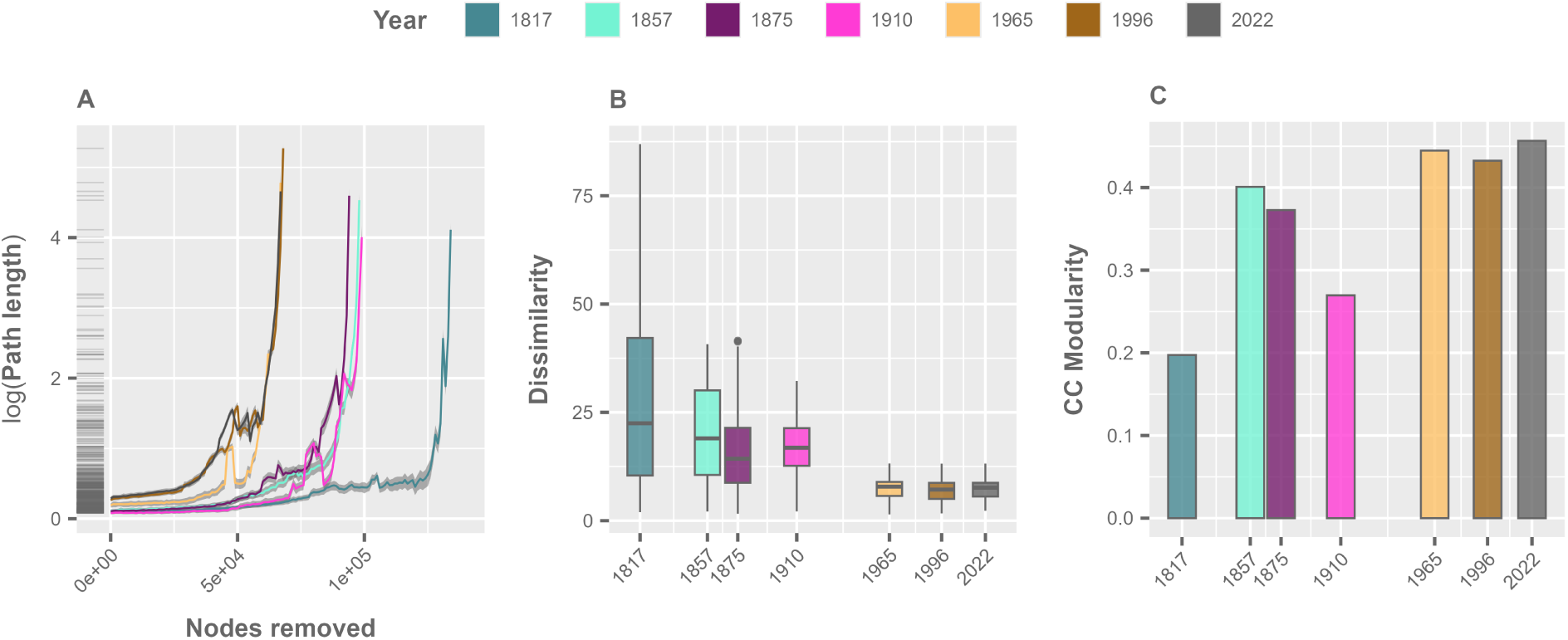
**A** Average reachability adjusted path lengths (*d^net^*) from 100 randomly sampled source nodes to all other reachable nodes (100 replicates per year; lines indicate the arithmetic mean and the standard deviation across replicates). Path length is plotted for different years and as a function of the number of nodes removed with increasing importance. **B** Dissimilarity of RCIs between layers of the multilayer network at different years. **C** Connectivity cluster (CC) modularity at different years.

The dissimilarity between network layers based on RCIs decreased substantially over time (figure 7B). While affecting both the average and variance, this development persisted from 1817 to 1965 before stagnating at a very low level. An effect of restoration measures was not indicated by layer dissimilarities. Compared to 1817 (system-wide CC1 dominance), CC modularity increased substantially until 1857, following a shift from CC1 to CC2. Until 1910, modularity declined but remained higher than in 1817 due to the emergence of two large spatially coherent areas upstream and downstream of Vienna, dominated by CC2 and CC4, respectively. By 1965, further habitat losses and continued river engineering led to greater lateral separation of different CCs and increased their connectivity longitudinally (along the general direction of flow). As a result, modularity increased again to a high level, followed only by minor changes until 2022.

## 4 Discussion and conclusion

In 1817, the investigated section of the Danube was surrounded by diverse and extensive floodplains and functioned as a highly connected and dynamic meta-ecosystem. Frequent hydrological exchange created a dense web of functionally highly connected and abundant habitats with high niche diversity, reflected by the predominance of CC1, low CC modularity, and strong cross-layer dissimilarity in the functional multilayer network, a pattern found today only in small subsystems (Recinos Brizuela et al., 2024). At this time, source–sink dynamics, such as rescue effects (Granzotti et al., 2021) and spatial subsidies (Junk et al., 1989), operated across life stages and functional groups (fish, amphibians, flying macroinvertebrates, non-flying macroinvertebrates, and plants). Thus, demographic risk was effectively distributed across space and time, ultimately stabilizing local and regional population and community dynamics. This strong system-level resilience was further underlined by the high structural robustness of the functional multilayer network.

Over the following two centuries, river engineering reorganized this baseline into a system with approximately 50% loss of aquatic habitat area, a shift away from the extensively anabranching structure, lotic waters, and generally dynamic conditions described by CC1. While river regulation measures were applied only locally until 1857, major engineering works, particularly within and around Vienna, were completed by 1875, followed by a system-wide implementation across the remaining area by 1910. While this transformation was already profound in the eastern part of the investigated system at the time, extensive levee construction extended the level of alteration also towards the western part (upstream of Vienna) by 1965. Subsequently, the construction of four major hydropower plants (three in the main channel) further altered functional continuity patterns through impoundment and continued floodplain disconnection until 1996, before the onset of ongoing restoration efforts. As a result of this long-term development, reaches located closer to the main channel or in tributaries, if not lost entirely, indicated a shift from CC1 to CC2 (side channels connected permanently only downstream; impounded main channel reaches) at first, and later from CC2 to CC3 (tributaries; non-impounded reaches of the main channel and major side channels, both with limited connection to the floodplain). Most reaches of the main channel shifted over time directly from CC1 to CC3, and following impoundment to CC2. Hence, the taxa associated with CC1 (SC1 and SC2) are likely to have been most affected over time. Besides several flying and non-flying macroinvertebrates and some plants, this group includes a very substantial share of the system’s native fish species (figure 10; appendix). Particularly, medium-distance migratory fish suffer from restricted access to dynamic sites located in tributaries, side channels, or structurally rich shorelines of the main channel, which are suitable for spawning and juvenile development (Pelz et al., 2026), leading to limited reproductive success and increased pre-recruitment mortality. Fragmentation of populations and local isolation further erode portfolio effects (Schindler et al., 2015), stemming from variations in spawning destination, timing, distance, and homing fidelity. As a consequence, vulnerability is heightened by concentrating risk among fewer locations and pathways. The negative effect of this development on MDM taxa, such as the Danube Salmon (*Hucho hucho*), is partly reflected by historical records from the Viennese fish market, where this iconic species disappeared at the beginning of the 20*^th^* century (Jungwirth et al., 2014). Furthermore, reaches classified as CC1 in 1817 have experienced an intense colonization by invasive macroinvertebrate and fish taxa (Liška et al., 2021). While introduced mainly through human activities, the extent to which invasive taxa within Gobiidae, Bivalves, and Crustacea dominate communities today further outlines the habitat degradation and the consequential loss of resilience.

Another major development over time was the increasing shift of peripheral reaches to CC4 (SC3 and SC4; isolated habitats). While these habitats became more abundant (at the expense of other CCs), the results already indicated ongoing losses since 1996 due to decreased hydrological connectivity, initiating terrestrialization processes. Nevertheless, taxa associated with less dynamic or isolated habitats (SC3 and SC4, respectively) experienced a manifold increase in habitat availability compared to the 1817 reference. However, relative to the increased amount of available habitat, functional connectivity decreased, potentially elevating local vulnerability. However, ongoing terrestrialization processes pose a substantially more pressing threat (Hohensinner et al., 2011). While this issue is generally caused by insufficient hydrological exchange, the root cause differs between impounded and free-flowing sections of the main channel. In the former case, where large parts of the floodplain are permanently submerged and hence lost, hydraulic exchange with the remaining parts is limited by regulated water levels or by levees. In free-flowing sections, the underlying cause is riverbed incision due to excess transport capacity, which in turn is caused by engineered channelization and by sediment loads being trapped upstream of transversal barriers (Surian and Rinaldi, 2003). As a result, water levels decline relative to average ground level, effectively disconnecting floodplains from the main channel, a development exacerbated by climate change (Klasz and Baumgartner, 2024). Hence, further habitat losses and a continued decline in functional connectivity are imminent without the implementation of appropriate mitigation or restoration measures. While the community adapted to more dynamic habitats, in particular migratory fish (MDM), has already been fundamentally decreased, the ongoing development leads to a scenario where also the community adapted to CC4 or even more isolated habitats (not included in this study), is likely to be affected to a similar or worse extent (Hamer et al., 2023; Pataki et al., 2013).

On a system-level perspective, the loss of functional connectivity severed critical routes for migration, life-stage transitions, and recolonization, thereby increasing isolation, demographic variance, and, ultimately, local extinction risks (Ahmad et al., 2025). This structural degradation is further reflected by the collapse of cross-layer dissimilarity (loss of functional diversity; Meng et al., 2022) and by increased spatial modularization of CC (functional isolation; Fletcher et al., 2013). While increased modularization can lead to more efficient buffering of disturbances (Gilarranz et al., 2017), the negative effects of functional isolation are likely to outweigh these benefits (Zhang et al., 2025). Accordingly, the system-level resilience to disturbances has likely declined drastically, as indicated by the decreased robustness of multilayer networks to node removal (Artime et al., 2024). Furthermore, the development observed in the investigated system must be viewed in the context of catchment-scale alterations. In particular, losses in longitudinal connectivity along the dendritic river network due to the construction of transversal barriers will have effectively further reduced the functional connectivity for MDM fish (Kowal et al., 2024, 2025).

River floodplain systems possess significant conservation and restoration potential by acting as biodiversity hotspots, refugia, stepping stones, and migration corridors at larger spatial scales (Funk et al., 2026; Robinson et al., 2002; Stoffers et al., 2024). These systems are therefore highly relevant for fulfilling the requirements of legislative frameworks such as the Nature Restoration Regulation (European Commission, 2024), the Water Framework Directive (European Commission, 2000), and the Habitats Directive (European Council, 1992). However, for the investigated system and many others, accelerating and scaling up restoration efforts is imperative to halt ongoing degradation (Grizzetti et al., 2017). Reestablishing dynamic habitat conditions and functional connectivity while halting terrestrialization can be achieved by reconnecting side channels already at low water levels and removing bank reinforcements. In addition, sediment transport across transversal barriers needs to be restored (Habersack et al., 2016). Otherwise, the riverbed incision in non-impounded reaches will continue to increasingly disconnect floodplains from the main channel, making any measures to restore hydrological connectivity and, thus, dynamic habitat conditions unsustainable in the long term. If technical solutions fail to restore sediment transport across barriers, there are ultimately only two management options: accepting the near-complete loss of what remains of a once extensive floodplain meta-ecosystem, or removing barriers where necessary. In any case, under a business-as-usual trajectory, system-level degradation and loss of resilience, intensified by climate change, will continue. Furthermore, functional connectivity for medium-distance migratory fish taxa requires the restoration of key dynamic habitats and the reestablishment of passability across transversal barriers (Seliger and Zeiringer, 2018). This can be achieved in part by installing large-scale, nature-like bypass channels that recreate some of the dynamic, lotic habitats lost to impoundment, or by updating existing fish migration aids if efficient passability has not yet been established. Lastly, continuous monitoring of ecological, hydrological, and geomorphological changes is necessary to verify the effectiveness of restoration strategies and adapt where necessary.

While the present study provides a novel, detailed, and long-term analysis of how river engineering has changed the functionality and resilience of a fluvial meta-ecosystem, the applied modeling framework introduces uncertainties that warrant consideration. Firstly, historical reconstructions validated primarily by contemporary observations limit direct inference about past and future population sizes. Secondly, network-based estimation of functional connectivity may overestimate realized connections where, e.g., varying hydraulic conditions or behavioral constraints additionally limit exchange. To better quantify population resilience of key species under climate change and management scenarios, future research may apply individual-based population models within a network representation of the investigated system to incorporate stochasticity, trait variation, and life-stage dependence of movement and survival. Embedding such an approach within a socioecohydrological framework will help identify feasible management scenarios with a higher likelihood of achieving long-term population persistence and, ultimately, system-level functionality and resilience.

In sum, two centuries of habitat loss, alteration, fragmentation, and homogenization have fundamentally reorganized and, in many ways, degraded the investigated system. As a result, populations will have declined substantially in size and face increased extinction risk due to diminished ability to buffer disturbances. Restoration measures implemented so far have not achieved system-level rehabilitation. The results point to an urgent need for large-scale restoration and measures to halt channel incision in order to restore dynamic floodplain habitats and prevent ongoing terrestrialization. Furthermore, the passability for fish and other aquatic organisms across transversal barriers needs to be effectively restored.

## Declarations

### Author Contributions (CRediT)

Johannes L. Kowal: Data curation, Formal analysis, Investigation, Methodology, Validation, Visualization, Writing - Original Draft, Writing - Review & Editing; Sarah M. Gross: Data curation; Gertrud Haidvogl: Data curation, Writing - Review & Editing; Thomas Hein: Conceptualization, Funding acquisition, Project administration, Writing - Review & Editing; Severin Hohensinner: Data curation, Writing - Review & Editing; Andrea Funk: Conceptualization, Data curation, Methodology, Supervision, Writing - Review & Editing.

### Data availability statement

The code and datasets are available online at https://doi.org/10.5281/zenodo.21165311.

## Appendix

**Figure 8:**
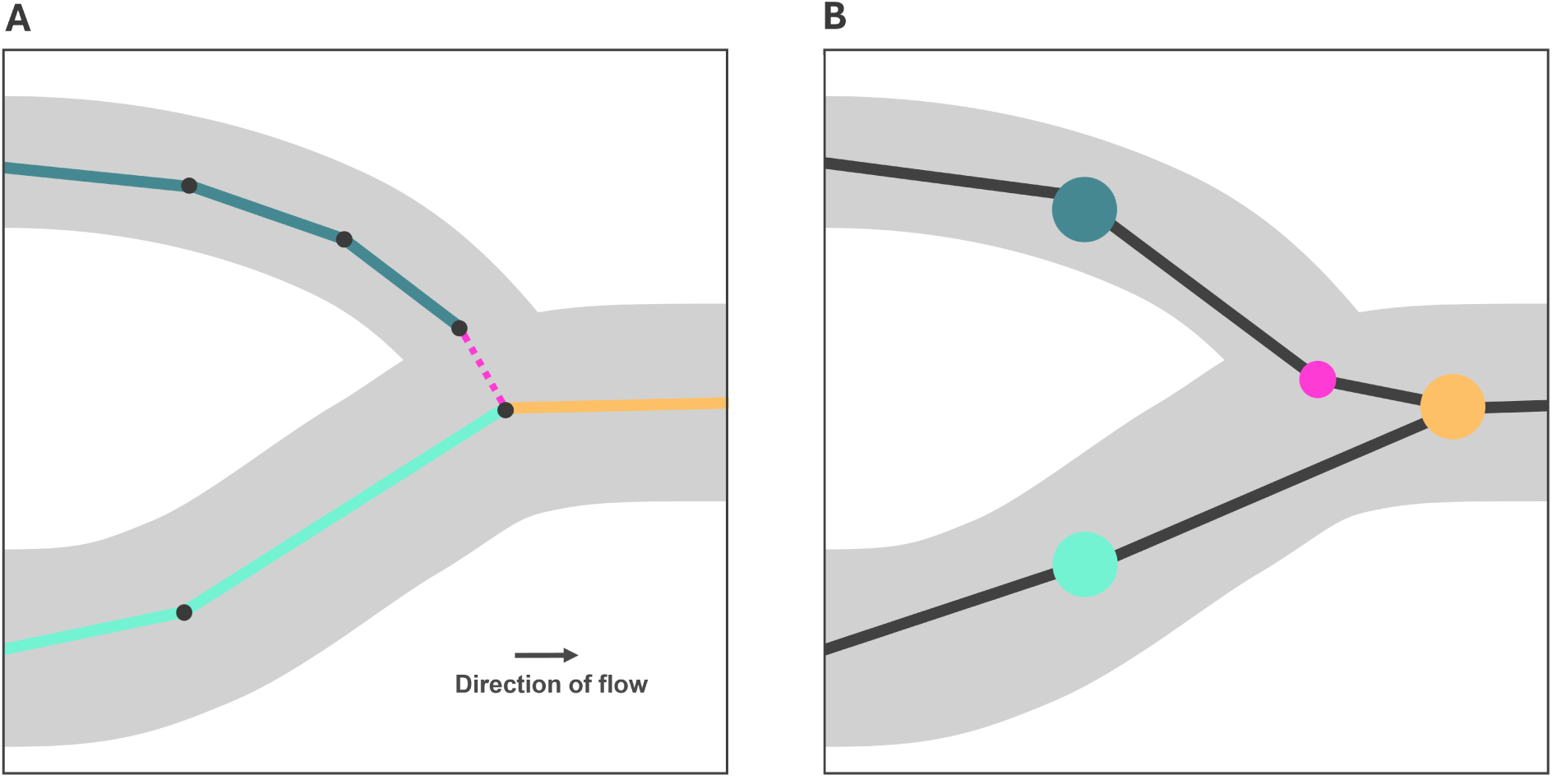
Illustration of a river confluence. **8A** Solid connected lines of the same color illustrate spatial line features representing waterbodies (reaches). Dotted lines represent connector line features. **8B** Illustration of the corresponding network. Large nodes reference line features of the corresponding color in figure 8A while small nodes reference connectors. Lines reference links in the direction of flow.

**Figure 9:**
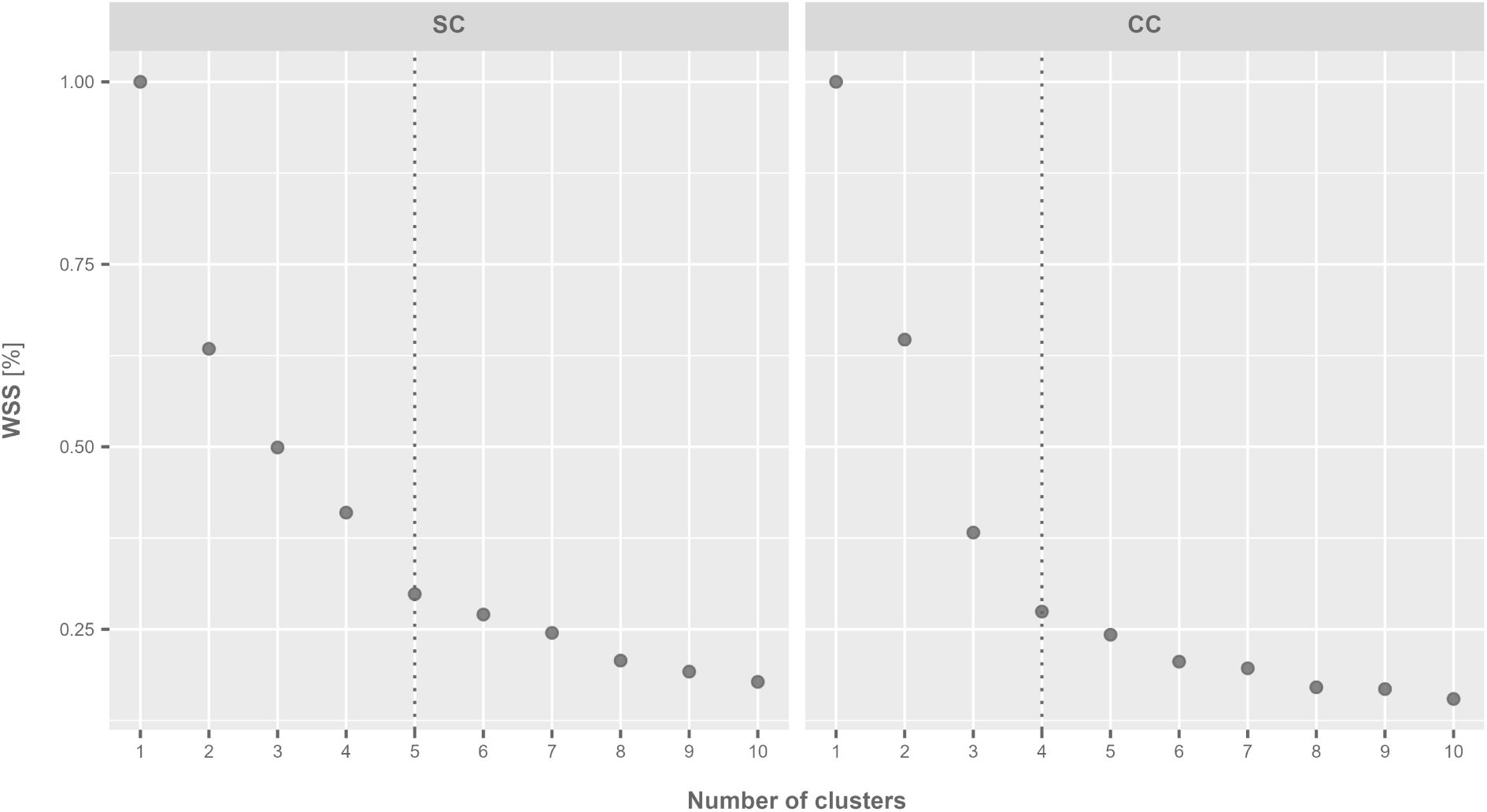
Within sum of squares (WSS) plotted against the number of clusters. The vertical dotted line indicates the final number of suitability (SCs) and connectivity clusters (CCs).

**Figure 10:**
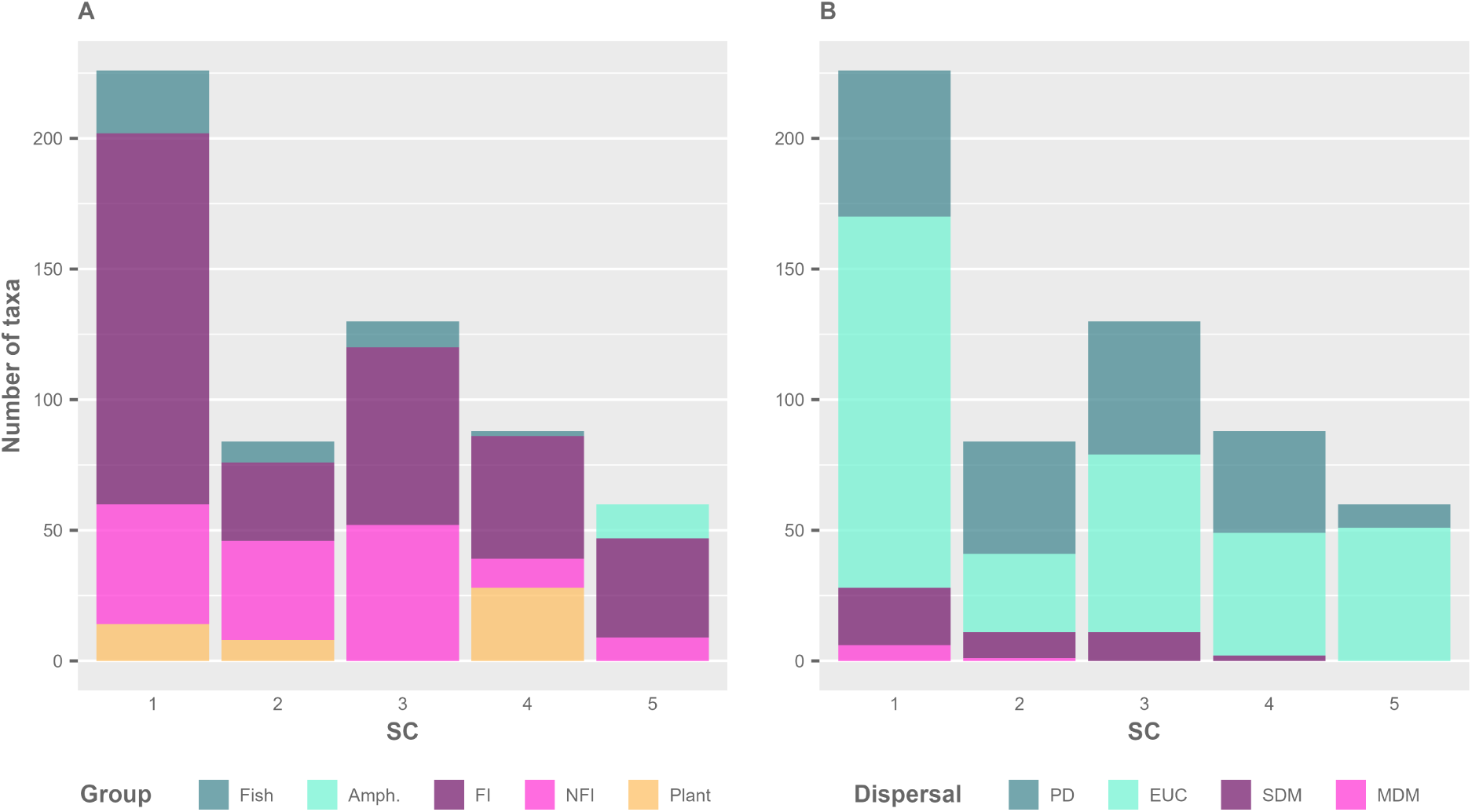
**10A**: Number of taxa per functional group and suitability cluster (SC). Functional groups are fish (Fish), amphibians (Amph.), flying macroinvertebrates (FI), non-flying macroinvertebrates (NFI), and plants (Plant). **10B**: Number of taxa per dispersal mode and SC. Dispersal modes are medium-distance migrators (MDM), short-distance migrators (SDM), passive dispersers (PD), and taxa dispersing more independently of the network structure (EUC).

**Figure 11:**
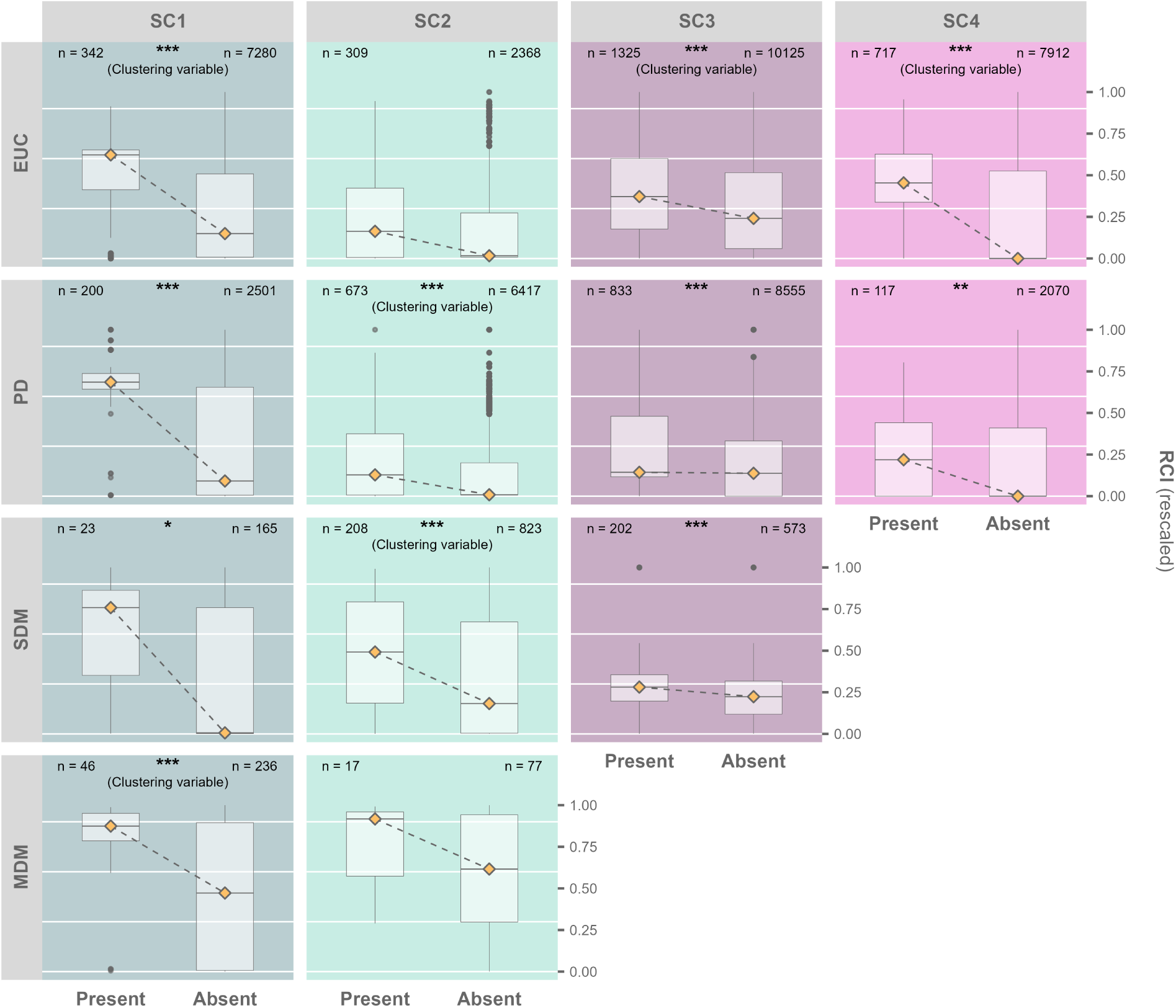
Reach connectivity index (RCI) configuration corresponding to combinations of dispersal modes and suitability clusters (SCs). “Present” and “Absent” group RCIs according to whether or not species associated with a given configuration were observed in a corresponding reach. The full data set is visualized, but significances were calculated based on a subset, randomly downsampled to an equal number of observations between “Present” and “Absent” (*N* indicates the number of samples in both groups before downsampling). Significance levels are indicated by asterisks (* *p <* 0.05, ** *p <* 0.01, *** *p <* 0.001), and variables used for the definition of connectivity clusters are labeled with “Clustering variable”.

**Table 2:** Data sources used for the mapping of reaches at different time steps. For additional information see Hohensinner (2025)

| Year | Source | Type | Scale |
| --- | --- | --- | --- |
| 1817 | Lorenzo (B II 82/B II 86)<br>Lower Austrian Provincial Library | Map | 1:28'800 |
| 1857 | Pasetti (B IX b 138)<br>Austrian Military Archives | Map | 1:28'800 |
| 1875 | 3<sup>rd</sup> Franzisco-Josephinian Land Survey<br>Austrian Military Archives | Map | 1:25'000 |
| 1910 | Map of the Austrian Danube<br>Private Archive Baumgartner | Map | Multiple |
| 1965 | Austrian Basemap (I)<br>Austrian Federal Office of Metrology and Surveying | Map | 1:50'000 |
| 1996 | Austrian Basemap (II)<br>Austrian Federal Office of Metrology and Surveying | Map | 1:50'000 |
| 2022 | World Imagery Basemap<br>ESRI Inc. | Satellite images | Multiple |

**Table 3:** Habitat suitability parameter *S_sc,h_* for different suitability clusters *sc* and habitats *h*.

| Suitability cluster | H1 | H2 | H3 | H4 | H5 |
| --- | --- | --- | --- | --- | --- |
| SC1 | 10 | 2 | 0 | 0 | 0 |
| SC2 | 1 | 6 | 2 | 0 | 0 |
| SC3 | 0 | 2 | 4 | 3 | 0 |
| SC4 | 0 | 0 | 3 | 8 | 0 |
| SC5 | 0 | 0 | 1 | 3 | 5.5 |

**Table 4:** Translation of habitats classified by Hohensinner et al. (2011) into habitats classified by the Chovanec et al. (2005).

| Habitat (Hohensinner et al., 2011) | Habitat (Chovanec et al., 2005) | Impounded |
| --- | --- | --- |
| Eupotamon A | H1 | No |
| Eupotamon B | H1 | No |
| Parapotamon A | (H1+H2) 0.5 | No |
| Parapotamon B | H2 | No |
| Plesiopotamon | (H3+H4) 0.5 | No |
| Tributary | H1 | No |
| Artificial | H4 | No |
| Eupotamon A | H2 | Yes |
| Eupotamon B | H2 | Yes |
| Parapotamon A | H2 | Yes |
| Parapotamon B | (H3+H4) 0.5 | Yes |
| Plesiopotamon | (H3+H4) 0.5 | Yes |
| Tributary | H2 | Yes |
| Artificial | H4 | Yes |

